# DNA-Based Nanoprobes for Fluorescence K+ Sensing in Neural Systems

**DOI:** 10.64898/2026.04.21.719852

**Authors:** Bryce Dunn, Sally Farag, Madinah Azizi, Laura McAuliffe, John R. Cressman, Remi Veneziano, Parag V. Chitnis

**Author notes:** These authors contributed equally to this work (Sally Farag, Madinah Azizi). First Author (Bryce Dunn).

## Abstract

**Significance:** Abnormalities in potassium ion concentrations across subregions of the hippocampus have been implicated in seizures and other pathologies. Direct measurements of potassium ion concentrations are largely made using invasive electrodes, which do not allow for wide spatial coverage. This fluorescent nanoparticle potassium sensor enables direct visualization of potassium dynamics and represents a minimally invasive alternative to electrode-based methods.

**Aim:** Here, we present a DNA-based fluorescence nanoprobe capable of sensing relative concentrations of potassium ions within populations of neurons. We present its effectiveness in monitoring neuronal K^+^ dynamics in response to electrical stimulation ex vivo.

**Approach:** We used widefield fluorescence microscopy to monitor changes in fluorescence intensity in labeled brain tissue in response to electrical stimulation ex vivo.

**Results:** We found that our nanoprobe could be retained within the intracellular compartment and modulate in fluorescence intensity linearly in response to induced electrical current. Our K^+^ Sensor showed a fractional fluorescence change of approximately 1% per 10 mA of applied stimulation current in brain tissue. Optical spectroscopy confirmed the selectivity of the nanoprobes to potassium ions over other endogenous ions.

**Conclusions:** Our findings indicate that this nanoprobe can be used to detect more complex potassium dynamics implicated in various pathologies of the nervous system, such as migraines, seizures, and trauma.

## 1 Introduction

Potassium is the most abundant intracellular cation that contributes to many physiological processes. In neurons, potassium (K^+^) plays a crucial role in signal transmission by maintaining the resting membrane potential, shaping action potential generation, and contributing to the refractory period through membrane hyperpolarization [1], [2], [3], [4]. The neuronal membrane potential closely reflects the potassium equilibrium potential, which is largely determined by the ratio of intracellular to extracellular K^+^ concentrations. Potassium is maintained at a 30–40-fold higher concentration inside cells (∼150 mM) than in the extracellular space (3.5–5.0 mM), contributing to a resting membrane potential of approximately −70 to −80 mV [5], [6], [7], [8]. During action potentials, neurons release K^+^ into the extracellular space during the repolarization phase, leading to transient increases in extracellular K^+^ concentration [9]. Activation of K^+^ channels promotes membrane hyperpolarization, stabilizing the membrane potential, and suppressing excessive excitatory signaling in neurons [10]. Disruption of K^+^ homeostasis alters neuronal excitability and synaptic transmission, and is associated with neurological disorders including epilepsy, Parkinson’s disease, autism spectrum disorder, and Alzheimer’s disease [11], [12], [13], [14], [15], [16]. Thus, elucidating K^+^ ion flux dynamics is essential for understanding pathological mechanisms driving these conditions.

Several approaches have been developed to monitor potassium dynamics *in vitro* and *in vivo*. Ion-selective electrodes have been a gold standard method for quantitative measurements of absolute potassium concentration [3], [17], [18], [19], [20]. While offering good temporal resolution, they are invasive and lack wide spatial coverage, constraining their use for monitoring spatiotemporal dynamics of K^+^ flux. Fluorescent indicators offer a complementary approach, enabling wide-field spatial coverage with high temporal resolution but without the tissue disruption associated with electrode placement. Most available fluorescent K^+^ sensors have been developed for extracellular measurements, with only a limited number designed for intracellular imaging [21], [22], [23], [24].

We have developed a fluorescent DNA nanoparticle-based sensor designed for the detection of potassium ions in biological environments. Our nanoprobe leverages a K^+^ aptamer, functionalized with a rhodamine derivative, ATTO Rho101 (ATTO-TEC, Ex. 587 nm, Em. 609 nm), and a quencher dye, IowaBlack (Integrated DNA Technologies, peak absorbance: 656 nm) with overlapping emission and absorbance spectra, respectively (Fig. 1a). Upon interacting with potassium ions, a conformational change in the DNA nanostructure results in G-quadruplex formation (Fig. 1b). The formation of the G-quadruplex brings the fluorophore and the quencher in close proximity, producing a FRET-based quenching effect [25], [26]. This real-time fluorescence quenching provides an optical measure of potassium ion concentration changes within the cellular milieu.

**Fig. 1.**
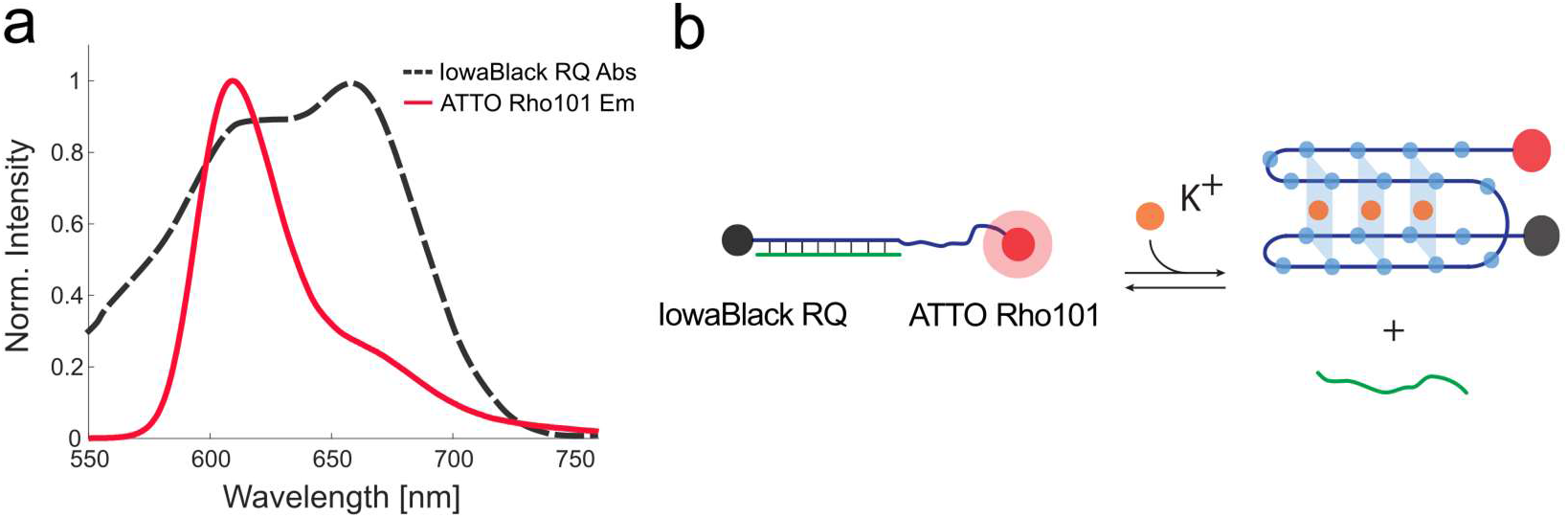
Optical components and molecular design of the DNA-aptamer potassium sensor. (a) Relative absorbance spectra of IowaBlack and relative emission spectra of ATTO Rho101. (b) Schematic of molecular design of the potassium sensor and FRET quenching during G-quadruplex formation.

In this study, we observed fluorescence intensity responses to intracellular potassium ion dynamics following electrical stimulation of the labeled mouse hippocampus. Timelapse imaging was performed to quantify changes in fluorescence intensity within the hippocampal neurons in response to induced electrical stimulation trains. Spectral measurements helped confirm selectivity and fluorescence response. These results are promising because intracellular K+ imaging is inherently challenging because the high resting potassium concentration (∼150 mM) necessitates sensing elements that must discriminate K^+^ from competing ions such as Na^+^. Our probe fills an unmet need for an intracellular, FRET-quenching fluorescent sensor for applications in neural tissue.

## 2 Materials and Methods

### 2.1 Synthesis of the K^+^ Sensor

The two DNA oligonucleotides strands (modified and non-modified) were purchased from Integrated DNA Technologies (IDT) in dried form, resuspended in water without further purification, and stored at -20ºC prior to being used. The sequences are listed in Supplementary Table S1. The K^+^ aptamer sensor strand is a 27 bases long single strand labeled in its 5’-end with an ATTO Rho101 dye and with Iowa Black RQ quencher at its 3’-end (Table S1). The K^+^ Sensor additionally includes a complementary blocker strand of 18 bases, which maintains the spatial separation of the dye and quencher by creating a duplex in the absence of potassium. The sensor was folded by combining both strands at an equimolar concentration of 10 µM in a TAE Folding Buffer (40 mM Tris, 20 mM acetic acid, 2 mM ethylenediaminetetraacetic acid [EDTA], and complemented with 12 mM magnesium chloride [MgCl_2_] at pH 8.0) with slow annealing over 2 hours from 90°C to 4°C in a Biorad T100 thermocycler. The sensor was then stored at 4°C until further use. The duplex formation was confirmed with polyacrylamide gel electrophoresis (PAGE), using a 20% gel in Tris acetate EDTA (TAE) buffer. 10 µL of samples at 5 µM were loaded with 1X Gel Loading Dye Purple (B7025S; New England BioLabs) and ran for 2 hours at 100 V. The gel was then stained for 10 minutes in a SYBR Safe (S33102; Thermo Fisher Scientific) solution and imaged with Azure c150 blue transilluminator.

### 2.2 Spectral Characterization of K^+^ Sensor

Measurements of fluorescence emission were carried out using a StarLine spectrometer (Avantes) and HAL-Mini halogen lamp (Avantes) and measured using AvaSoft (Version 8). Measurements were made in quartz cuvettes (ThorLabs) using 100 µL samples. Emission response to potassium ions was conducted in K^+^ Sensor diluted in milliQ water. Emission spectra were measured in a range from 500-700 nm using broadband excitation. Investigations of metal cation effect in the concentration range of 0-20 mM consisted of serial dilutions of a concentrated solution of 1 M KCl, 1 M NaCl, 1 M MgCl_2_, or 1 M CaCl_2_ (Sigma Aldrich), followed by stirring. Studies of the metal cation effect were carried out at 25°C.

### 2.3 Cellular Imaging

HeLa cells were cultured on an uncoated #1.5 glass coverslip in media for 24 hours. Samples were labeled with FITC at 500 nM, Hoechst 33342 (NucBlue), and K^+^ Sensor at 500 nM. Microscopy imaging experiments were performed with a Zeiss LSM800 inverted microscope stand. FITC channel acquired 493 nm excitation and 400–575 nm emission detection at 2.0 Airy units. K^+^ Sensor channel acquired with 587 nm excitation and 565–700 nm emission detection at 2.5 Airy units. Hoechst channel was acquired with 348 nm excitation and 400–600 nm emission detection at 2.0 Airy units. The laser scan pixel time for every channel is 1.469E-06 sec. Confocal fluorescence images were taken using a 63×/1.4 Plan Apochromat oil immersion objective. Voxel size: 0.0706×0.0706×0.37 μm^3^ with 16 bits per pixel.

### 2.4 Animal Tissue Imaging

Acute hippocampal slices (400 µm-thick) were prepared from Adult C57BL6/J mice (male and female, P30-90) using a tissue chopper. Slices were incubated at room temperature for one hour in prepared artificial cerebrospinal fluid (126 mM NaCl, 3 mM KCl, 2 mM CaCl_2_, 2 mM MgCl_2_, 2 mM NaH_2_PO_4_, 26 mM NaHCO_3_, 10 mM D-Glucose), aerated with carbogen, prior to labeling. For fixed imaging, brain slices were stained with 500 nM of K^+^ Sensor and mounted between a glass slide and a #1.5 glass coverslip with a plastic spacer using ProLong mounting medium. Imaging was performed using a Zeiss LSM800 inverted microscope stand. Widefield fluorescence image was taken using a 10×/0.25 NA Achroplan air objective and Axiocam 820 camera. K^+^ Sensor channel acquired with 587 nm excitation and 565–700 nm emission detection. Image size: 1227.52×1236.288 μm (1120×1128 pixels).

Live recordings of brain slices were performed as follows: brain slices were stained with 500 nM of K^+^ Sensor and mounted in a custom recording chamber for immersion imaging in prepared artificial cerebrospinal fluid perfused with carbogen (Roberts Oxygen, 95% O_2_, 5% CO_2_). Images were acquired on a BX51WI Upright Microscope (Olympus) custom-modified with a translational stage, configured for widefield epifluorescence microscopy and controlled with Micro-Manager (MMStudio Version 2.0.0), equipped with a 10×/0.3 LUMPlanFL water immersion objective (Olympus). Images had a width and height of 3200 pixels, 1 plane (z), and 1 channel. K^+^ sensor was excited with a LS200US Metal Halide light source (Lumen) set to 100%; wavelength selection was carried out using a FEBP532-30 bandpass filter (ThorLabs), DMLP605R dichroic mirror (ThorLabs), and a longpass emission filter FELH0600 (ThorLabs). Images were acquired on a Kinetix sCMOS camera (Teledyne) with an exposure time of 50 ms with 1×1 binning.

Electrical stimulation pulses were delivered using a Model 2100 Isolated Pulse Stimulator (A-M Systems). Voltage measurements were made using a Digidata 1440 Series digitizer (Molecular Devices) and Clampex software (Version 10).

### 2.5 MTT Assay for Cell Viability

The MTT assay for cell viability (Roche) was performed using a microplate reader (Thermo Fisher) running SkanIt (Version 7.0). HeLa cells were cultured and seeded into a 96-well plate at 5000 cells per well and incubated at 37°C for 24 hours. Cells were expected to reach approximately 10,000 per well at the time of treatment, based on standard doubling time. Treatment groups (n = 5) were administered dimethyl sulfoxide, and concentrations of K+ Sensor (250 nM, 500 nM and 1000 nM) directly into the well plate before adding 10 µL of MTT. The well plate incubated overnight as the MTT was converted to formazan. After this incubation period, the formazan was dissolved using a solubilization buffer containing sodium dodecyl sulfate (SDS). Optical absorbance was read at 570 nm. Wells filled with molecular biology grade water were used as blanks. Results (Fig. S2) were normalized to cells that were not incubated with the K^+^ Sensor or DMSO. DMSO was used as a separate, negative control.

## 3 Results

### 3.1 Synthesis and characterization of the K^+^ Sensor

To monitor K^+^, we designed an aptamer-based sensor (K^+^ aptasensor) using a modified version of the design proposed by Yang et al. [27]. Here we used the K^+^ aptamer strand labeled with ATTO Rho101 and IowaBlack RQ at its 5’- and 3’-ends, respectively and hybridized it with a complementary blocker strand (Figure 1b and table S1). The blocker strand creates a duplex to maintain the spatial separation of the fluorophore and the quencher in the absence of potassium ions. To validate the assembly of the duplex, we ran a 20% PAGE gel with 10 µL of each strand along with the duplex at 5 µM concentration seen in Figure S1. The results show a clear formation of the duplex with increased molecular weight and no byproduct as expected for this type of construct.

### 3.2 Spectral Characterization of K^+^ Sensor

We performed optical characterization to demonstrate two key design points: emission quenching in response to potassium ions, and selectivity of the sensor to endogenous metal ions as measured by fluorescence emission intensity. The response of the nanoprobe (100 nM in deionized water) to increasing potassium ion concentration, shown in Fig. 2a, demonstrates emission intensity decrease. This emission quenching is further characterized in Fig. 2b by showing a linear relationship between intensity at the peak emission wavelength (610 nm, Fig. 2b cutaway plot) and potassium ion concentration, which is conserved for varying concentrations of K^+^ Sensor. Fig. 2c demonstrates the selectivity of the K^+^ Sensor when tested against common endogenous metal ions. It is known that there are approximately 100 mM potassium ions in living cells, and when undergoing depolarization, the concentration may shift by approximately 10 mM [28], [29]. Together, these results indicate a potassium-selective, fluorescence emission intensity quenching response in physiologically relevant ion concentrations.

**Fig. 2.**
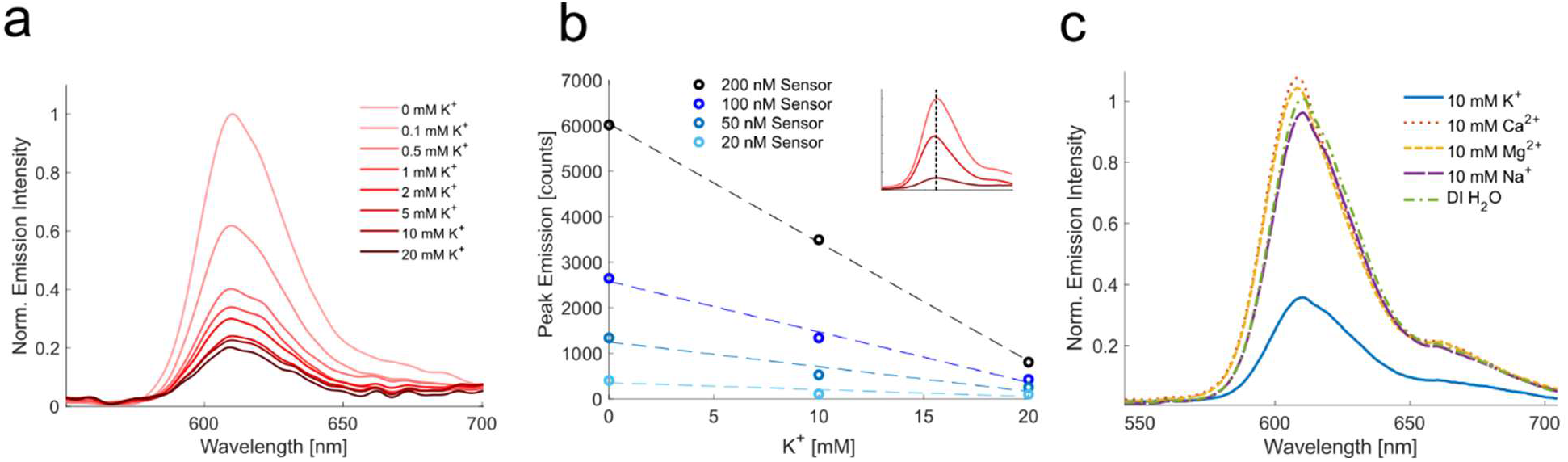
Optical properties of the DNA-aptamer potassium sensor in solution. (a) Relative emission intensity of 100 nM potassium sensor in 0, 0.1, 0.5, 1, 2, 5, 10, and 20 mM potassium chloride solution. (b) Peak emission intensity at 610 nm wavelength (cutaway) by potassium concentration for varying concentrations of potassium sensor. (c) Relative emission intensity of 100 nM potassium sensor in 10 mM potassium chloride, 10 mM calcium chloride, 10 mM magnesium chloride, 10 mM sodium chloride, and deionized water.

### 3.3 Cellular Imaging

We performed labeling and imaging experiments to show the spatial distribution of the K^+^ sensor in cell and tissue models. A HeLa model labeled with FITC, K^+^ Sensor, and Hoechst (Fig. 3a) demonstrates that the K^+^ Sensor enters the intracellular space between the plasma membrane and nuclear membrane. Labeling an acute brain slice with K^+^ Sensor shows cellular uptake of the nanoprobe into the bodies of hippocampal neurons (Fig. 3b). Additionally, the application of K^+^ Sensor in a HeLa model does not significantly impact cell viability or metabolic activity compared to untreated cells (0 mM K^+^ Sensor), as measured via MTT assay (Fig. S2).

**Fig. 3.**
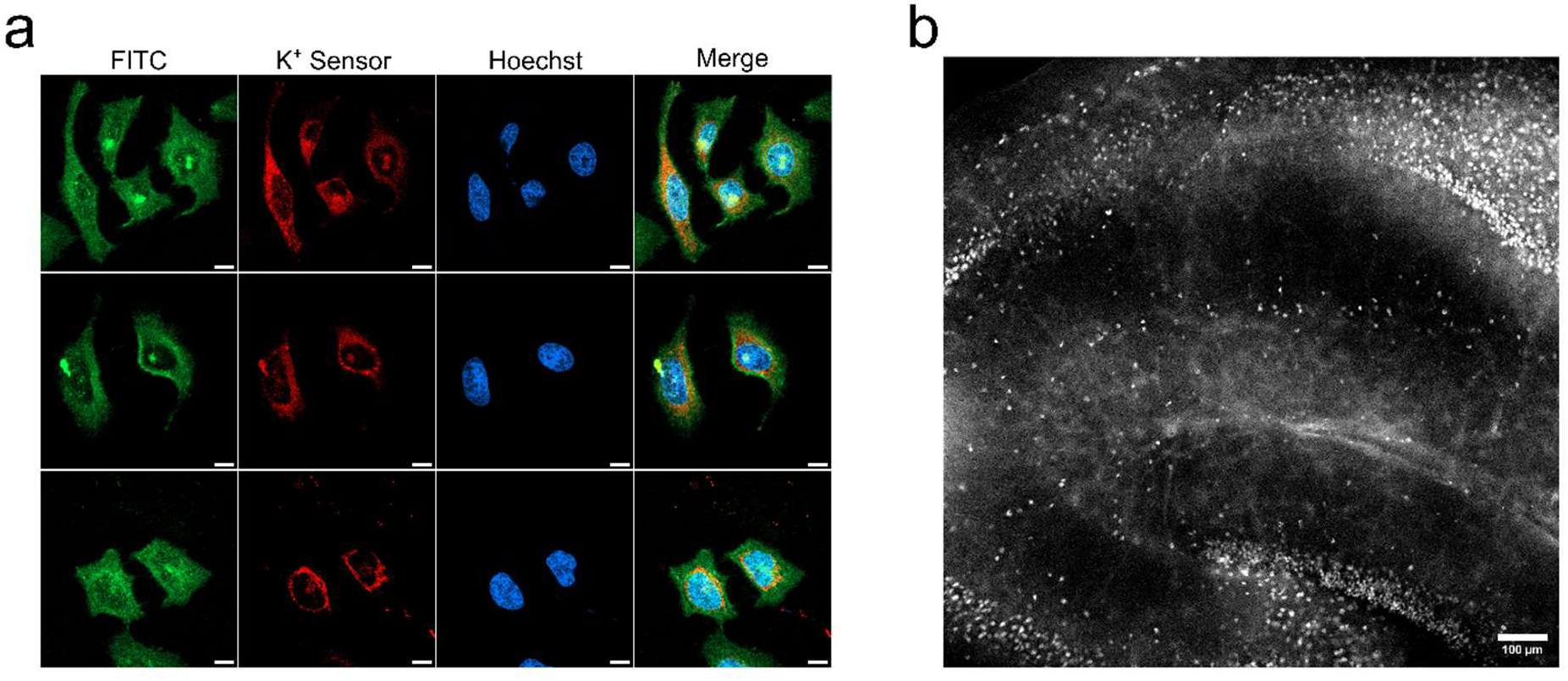
K^+^ sensor labeling of cell and tissue models. (a) Confocal fluorescence microscopy images of HeLa cells labeled with FITC (green), K^+^ Sensor (red), and Hoechst (blue). Confocal images are presented as maximum intensity projections. Scale bar: 10 μm. (b) Widefield fluorescence image of acute hippocampus slice labeled with K^+^ sensor. Scale bar: 100 μm.

Confocal microscopy images confirm that the nanoprobe is internalized within the cell, where it interacts with intracellular potassium ions. Several studies have produced extracellular nanoparticle potassium sensors, and few have produced intracellular sensors [24], [27]. A study by Müller et al. produced a BODIPY-based intracellular potassium ion sensor for neural applications [30]. Their results suggest that energy-dependent endocytosis is the primary mechanism of cell entry. We observed efficient labeling of acute brain slice tissue in thirty minutes, in contrast to three-hour staining observed in the study by the Müller group.

### 3.4 Electrical Stimulation of Murine Brain Tissue

After confirming intracellular K^+^ sensor labeling, we demonstrated fluorescence intensity changes in tissue following pulsed electrical stimulation in labeled mouse brain tissue. The aim of this experiment is to visualize electrical stimulation of the canonical Schaffer Collateral [31], [32], stimulating the tissue in the CA3 region of the hippocampus and recording fluorescence response in the CA1 region (Fig. 4a). Using electrical stimulation paradigms of 2.5, 20, 30, and 40 mA, corresponding fluorescence intensity responses are observed (Fig. 4b). Stimulation currents in the milliampere range were required to produce detectable population-level fluorescence responses, likely due to current losses at the extracellular electrode-tissue interface and the requirement for synchronous recruitment of labeled neurons to generate a measurable optical signal. Responses were not observed at microampere-range currents. Neuronal cell bodies labeled with K^+^ Sensor demonstrate fluorescence intensity changes of approximately 0.5% in response to 2.5 mA stimulation current (Fig. S3) where a recording electrode in the extracellular space shows a 12-mV potential change. Fluorescence response to higher, 40 mA electrical stimulations can be visualized proximally to the stimulating electrode (Fig. S4). There is variation between sensor responses depending on several factors such as the precise positioning of the stimulating electrode within the CA3, positioning of the recording electrode in CA1, and positioning of the region of interest for analysis.

**Fig. 4.**
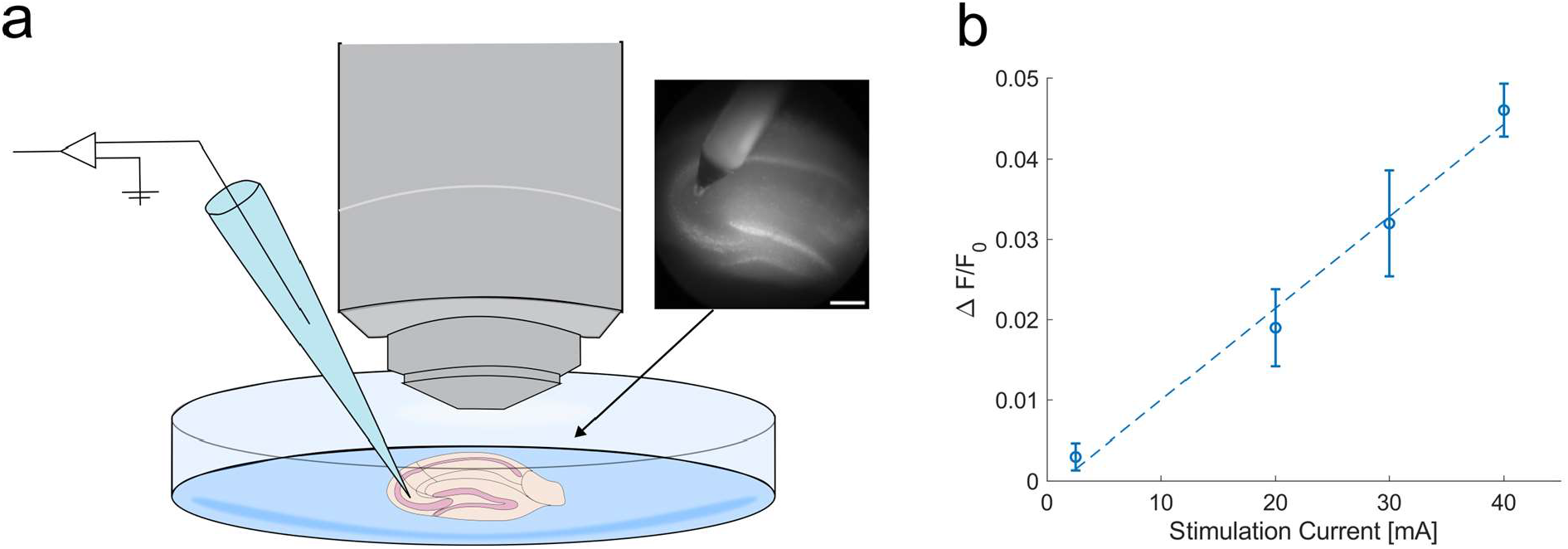
Electrical stimulation of labeled acute brain slices. (a) Schematic representation of sample and electrode placement in the recording chamber. Cutaway is a widefield fluorescence image of the labeled hippocampus with the stimulating electrode. Scale: 200 μm. (b) Relationship between fluorescence intensity changes of labeled brain tissue in response to electrical stimulation, by amperage of electrical pulse train. Linear trendline (dashed) is interpolated between data points (R^2^ = 0.9889).

Sensor response was maintained after repeated cycles of stimulation and higher amperages (Fig. S5). This is consistent with Moreira et al., who studied the effect of Iowa Black RQ on DNA duplex stability, which ranked among the most stabilizing quenchers in a comparison study [33]. The intracellular nature of the K^+^ Sensor is more relevant to both physiology and pathology, where the source of the K^+^ fluctuations is the tissue itself [22]. Our K^+^ Sensor showed a fractional fluorescence change of approximately 1% per 10 mA of applied stimulation current, which allows for the visualization of population-scale activity in live brain tissue.

## 4 Concluding Remarks

We have presented the design, synthesis, characterization, and biological applications of an aptamer-based potassium ion-sensing fluorescent nanoprobe. This nanosensor can monitor dynamic changes in intracellular K^+^ levels upon external stimulation. These findings demonstrate that this nanoprobe may serve as a tool for real-time imaging of potassium ion-dependent processes in neural tissue. This is a first-generation probe for imaging of intracellular K^+^ pools. However, the limitation of ∼1% fractional fluorescence change per 10 mA stimulation in tissue is notable and will be a subject of improvement in future iterations of the probe design.

### 4.1 Future Directions

Future studies will focus on interchanging the selection of fluorophore and quencher pairs, including those that emit in NIR wavelengths, and conjugating the platform with targeting ligands for cell-specific binding. ICG dye is a candidate for conjugation to the aptamer sequence, given that it emits in the NIR range and can penetrate living cells without inducing immediate degenerative changes [34]. Ionic content in the intracellular space is known to cause dye aggregation, quenching its fluorescence, although the effects of potassium have not been studied specifically [35]. Conjugation to the oligonucleotide scaffold may help prevent dimerization, but other techniques, such as lipid encapsulation, may also achieve this effect by shielding the dye from the ionic aqueous environment [36]. With these improvements, sensor performance may be evaluated in models of epileptic activity in tandem with EEG recordings. Future iterations of the probe design will be used to monitor and record potassium ion dynamics during seizure-like activity and in other model systems including cardiac models.

## Animal Ethics Statement

All animal procedures were approved by the Institutional Animal Care and Use Committee (IACUC) at George Mason University (Protocol #0532) and conducted in accordance with the National Institutes of Health Guide for the Care and Use of Laboratory Animals. Adult C57BL6/J mice (male and female, P30-90) were used for all experiments. Acute hippocampal brain slices were prepared as described in Section 2.4, and all efforts were made to minimize animal suffering and reduce the number of animals used.

## Code, Data, and Materials

Data are available from the corresponding authors upon reasonable request.

## Acknowledgments

This work was funded through the NSF award no. 2128821, NIH Project Number 1R01EB037718-01 and the Presidential Scholarship at George Mason University. The authors acknowledge the support of the IPN Microscopy Suite at George Mason University and the material contributions of the laboratory of Ted Dumas.

## Appendix A Supplemental Material

The following supplemental figures support the main text findings.

**Table 1.**
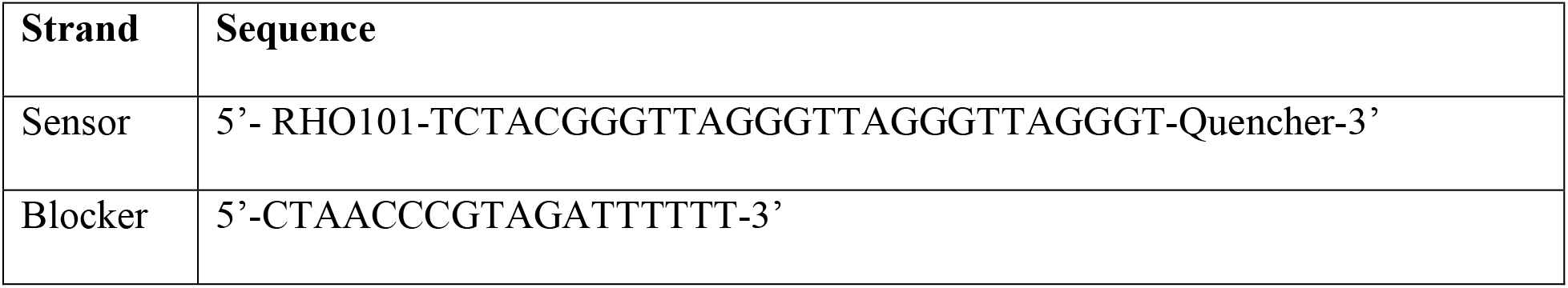
DNA sequences for K^+^ Sensor.

**Fig. S1.**
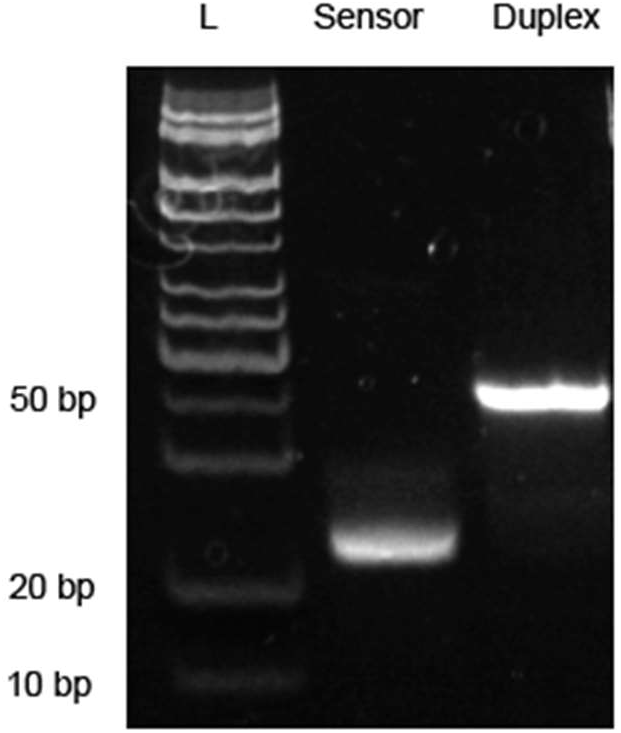
20% PAGE gel with the Ultra Low Range DNA Ladder, Sensor strand and assembled K^+^ Sensor Duplex from left to right.

**Fig. S2.**
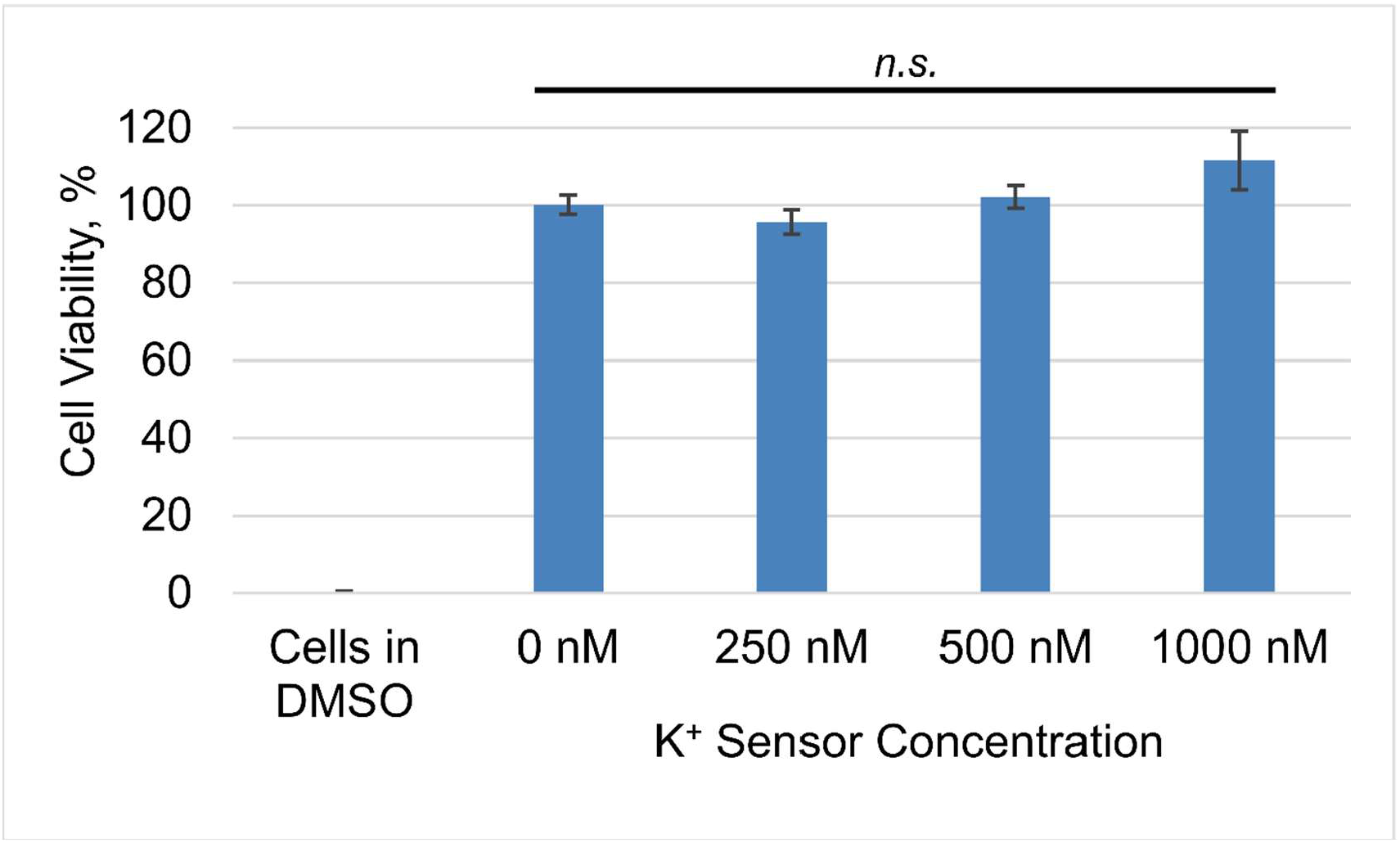
Cell viability MTT assay results. Data are presented as mean ± SE; n.s. indicates no statistical significance between group means.

**Table 2.**
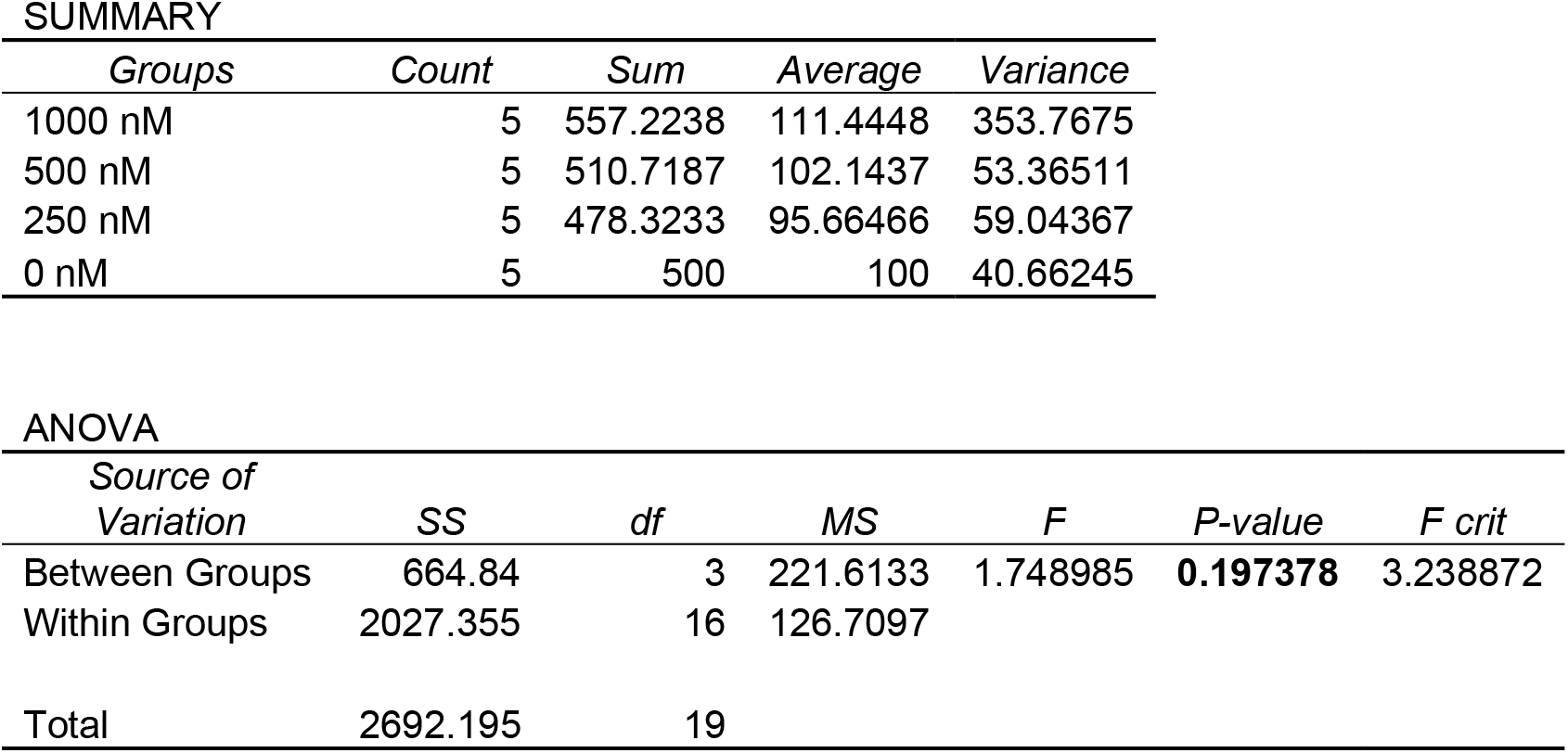
Single factor ANOVA results and group summary for K^+^ Sensor cell viability assay (H_0_: μ_1_ = μ_2_ = μ_3_ = μ_4_, α = 0.05).

| Groups | Count | Sum | Average | Variance |
| --- | --- | --- | --- | --- |
| 1000 nM | 5 | 557.2238 | 111.4448 | 353.7675 |
| 500 nM | 5 | 510.7187 | 102.1437 | 53.36511 |
| 250 nM | 5 | 478.3233 | 95.66466 | 59.04367 |
| 0 nM | 5 | 500 | 100 | 40.66245 |

**Fig. S3.**
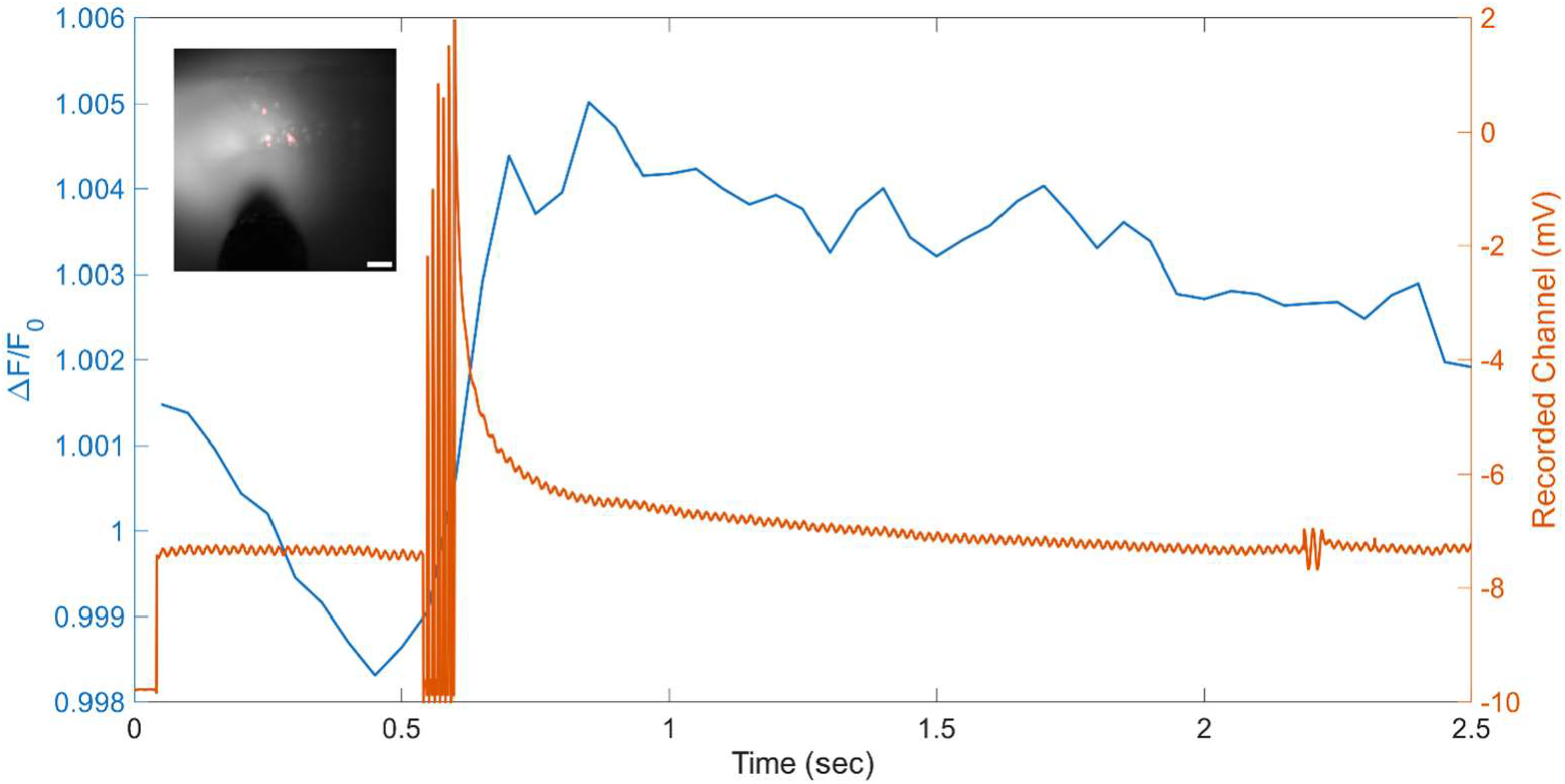
Fluorescence (blue) and voltage response (orange) to 2.5 mA electrical stimulation. Fluorescence response was recorded in CA3 cell bodies (cutaway, scale: 100 μm).

**Fig. S4.**
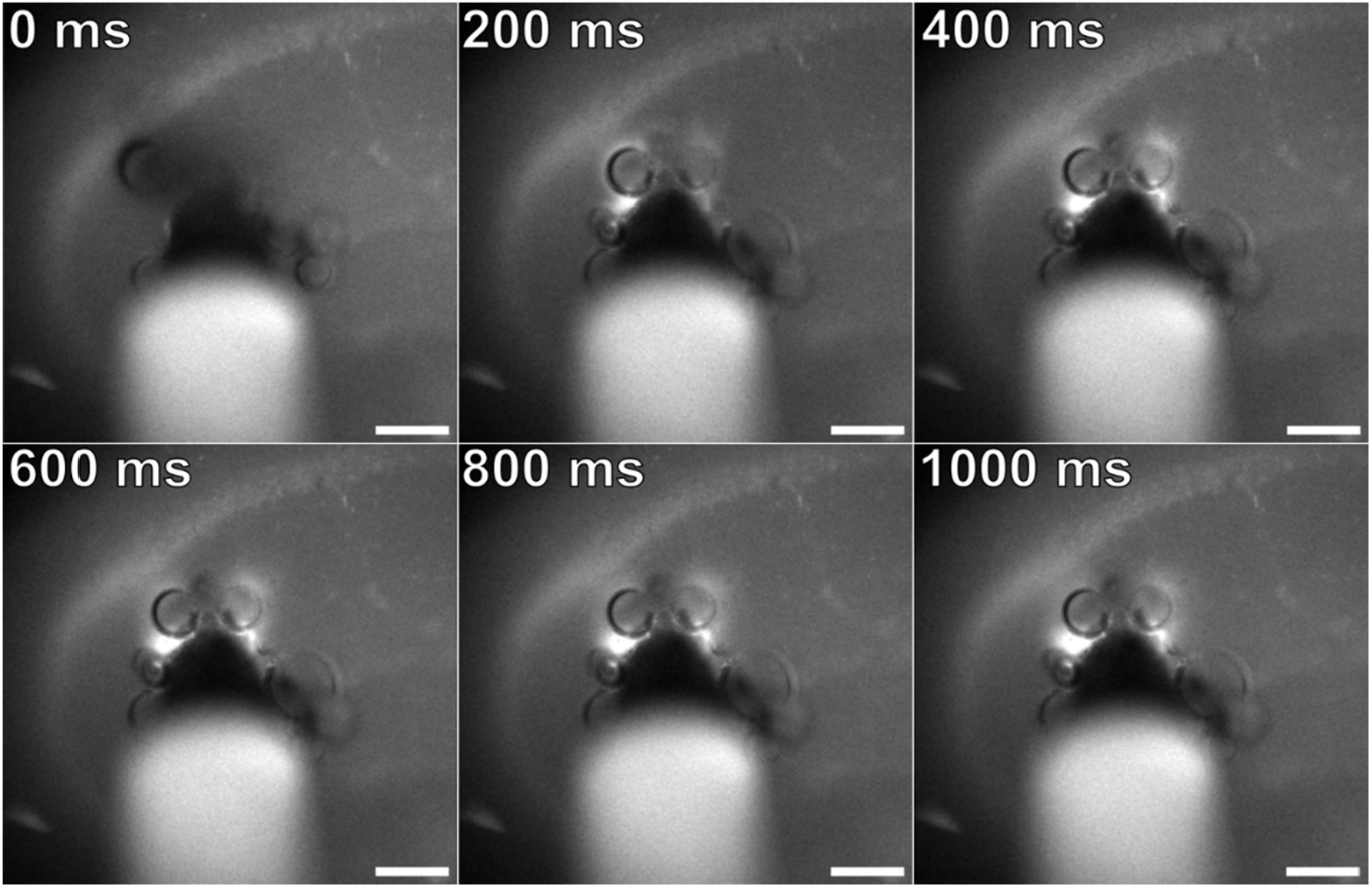
Timelapse montage of fluorescence response to 40 mA electrical stimulation in labeled mouse hippocampus (scale: 100 μm).

**Fig. S5.**
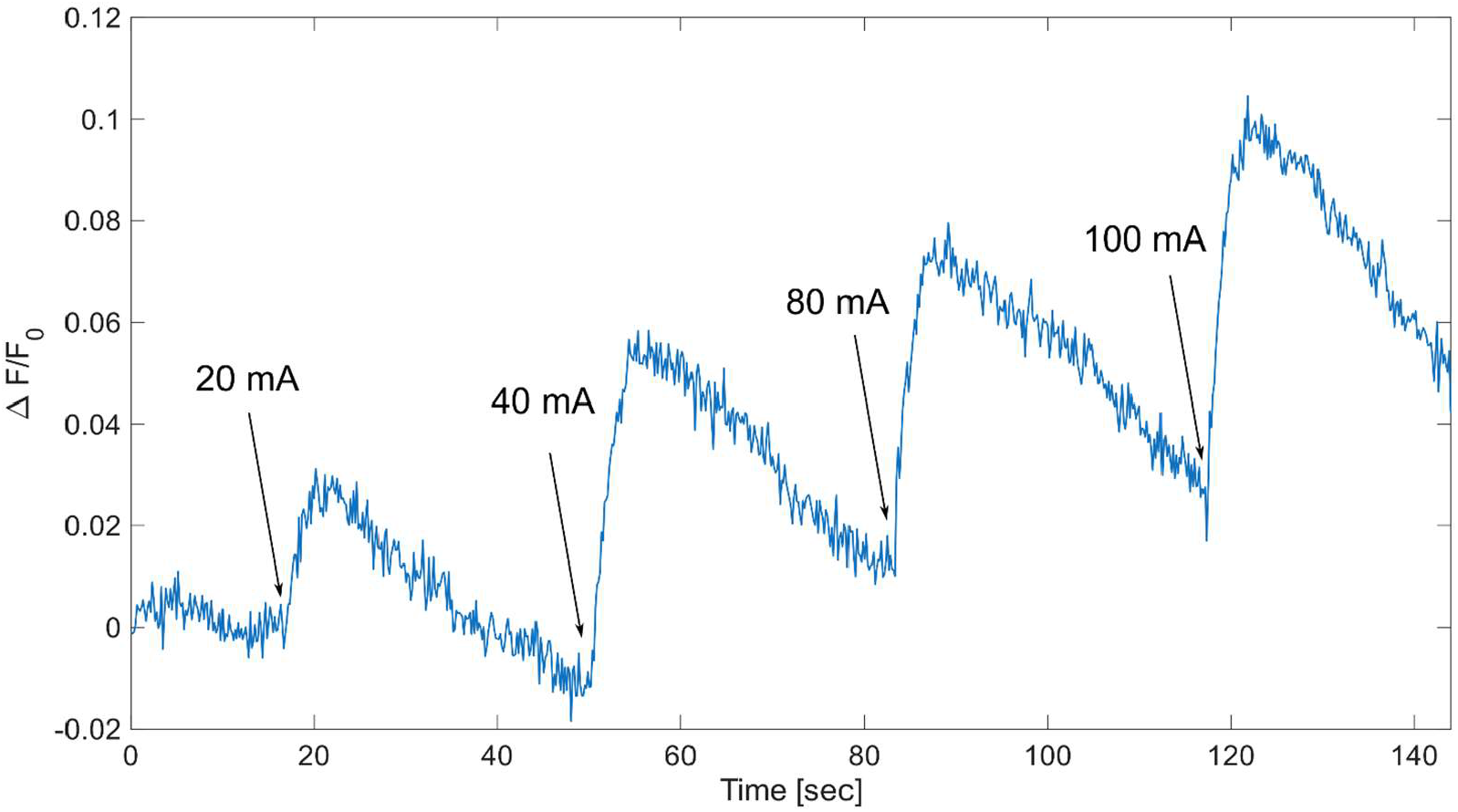
Fluorescence intensity responses to repeated electrical stimulations of progressively higher amperages.

