## Supplemental Figures for "DNA-Based Nanoprobes for Fluorescence K+ Sensing in Neural Systems"

### Appendix A Supplemental Material

The following supplemental figures support the main text findings.

**Table S1** DNA sequences for K^+^ Sensor

| **Strand** | **Sequence** |
| --- | --- |
| Sensor | 5’- RHO101-TCTACGGGTTAGGGTTAGGGTTAGGGT-Quencher-3’ |
| Blocker | 5’-CTAACCCGTAGATTTTTT-3’ |

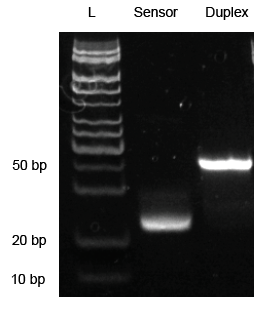

**Fig. S1** 20% PAGE gel with the Ultra Low Range DNA Ladder, Sensor strand and assembled K^+^ Sensor Duplex from left to right.

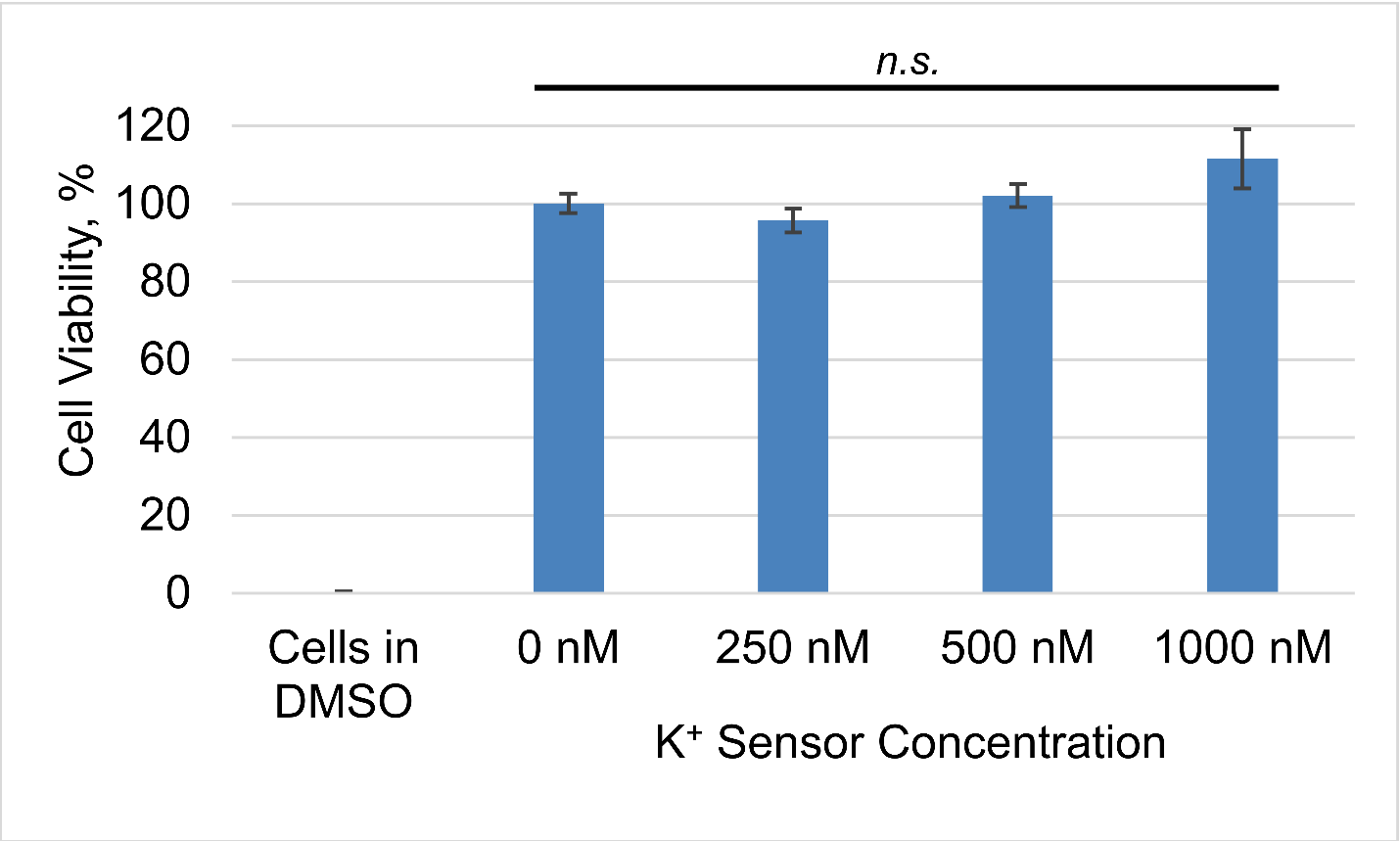

**Fig. S2** Cell viability MTT assay results. Data are presented as mean ± SE; n.s. indicates no statistical significance between group means.

**Table 2.** Single factor ANOVA results and group summary for K^+^ Sensor cell viability assay (H_0_: μ_1_ = μ_2_ = μ_3_ = μ_4_, α = 0.05).

| SUMMARY | |  |  |  |  |  |
| --- | --- | --- | --- | --- | --- | --- |
| *Groups* | *Count* | *Sum* | *Average* | *Variance* |  |  |
| 1000 nM | 5 | 557.2238 | 111.4448 | 353.7675 |  |  |
| 500 nM | 5 | 510.7187 | 102.1437 | 53.36511 |  |  |
| 250 nM | 5 | 478.3233 | 95.66466 | 59.04367 |  |  |
| 0 nM | 5 | 500 | 100 | 40.66245 |  |  |
| ANOVA |  |  |  |  |  |  |
| *Source of Variation* | *SS* | *df* | *MS* | *F* | *P-value* | *F crit* |
| Between Groups | 664.84 | 3 | 221.6133 | 1.748985 | **0.197378** | 3.238872 |
| Within Groups | 2027.355 | 16 | 126.7097 |  |  |  |
| Total | 2692.195 | 19 |  |  |  |  |

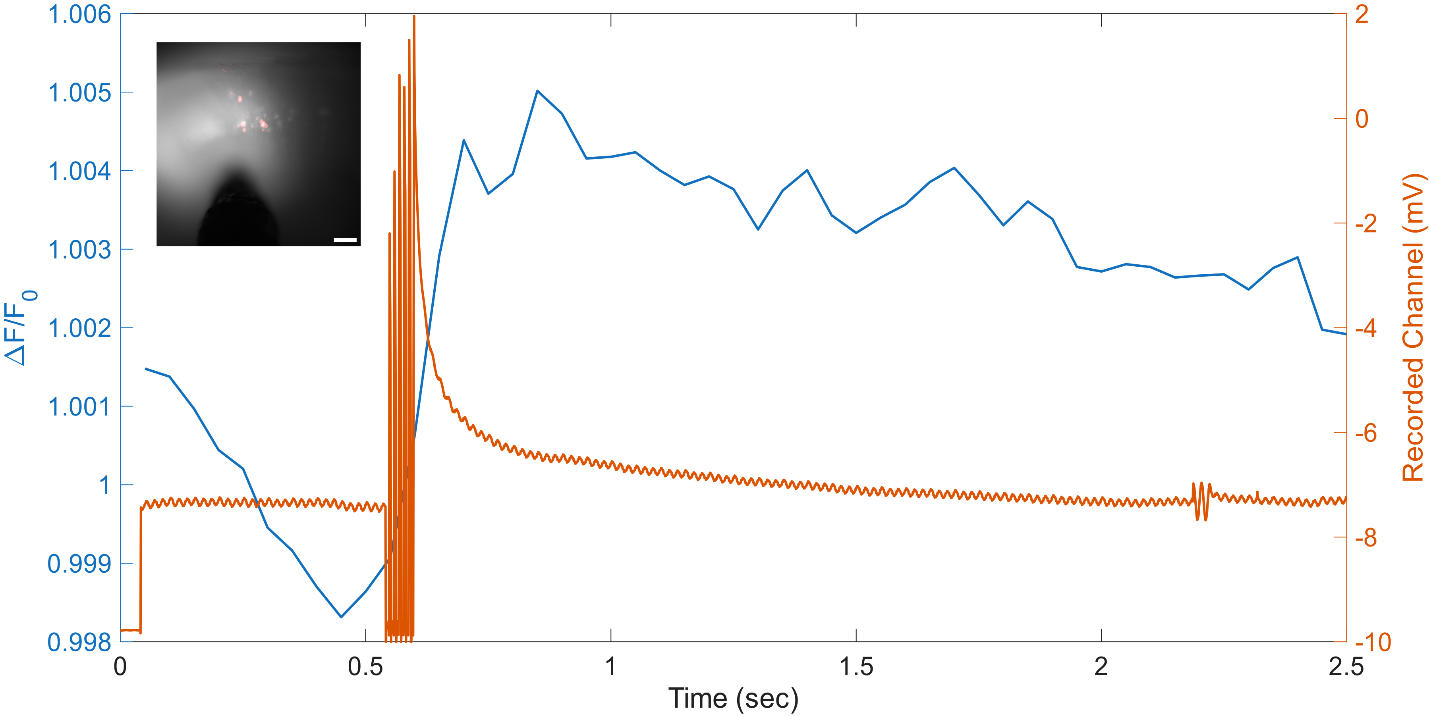

**Fig. S3** Fluorescence (blue) and voltage response (orange) to 2.5 mA electrical stimulation. Fluorescence response was recorded in CA3 cell bodies (cutaway, scale: 100 μm).

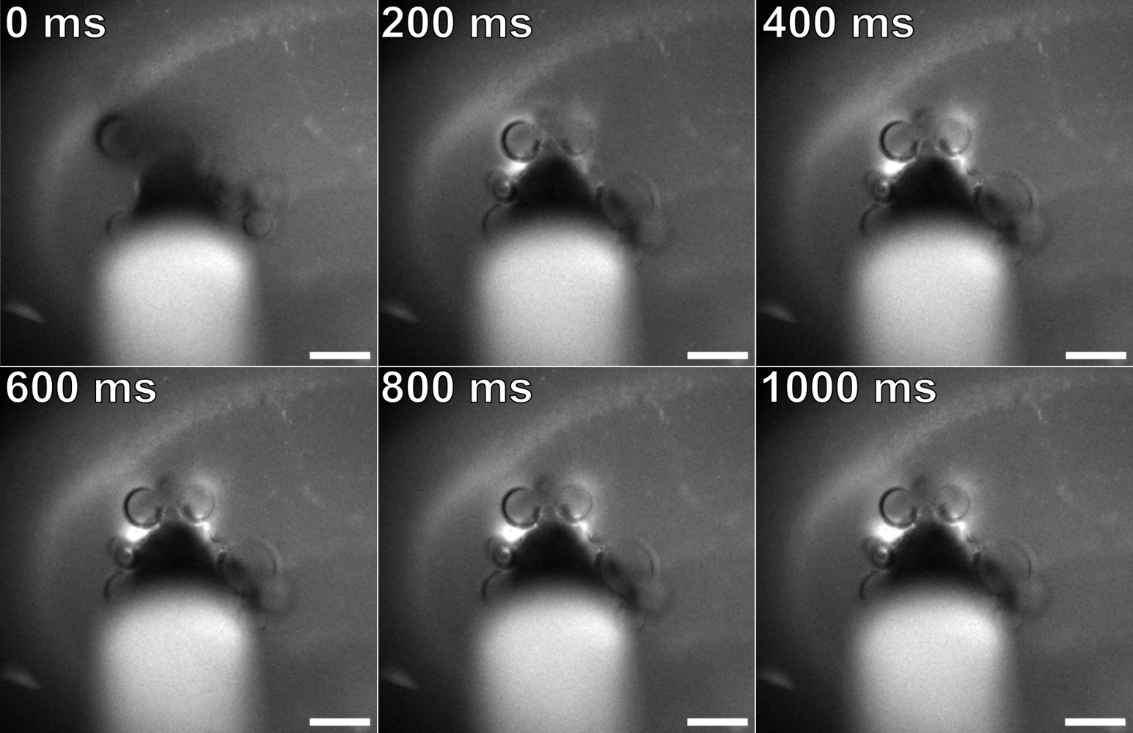

**Fig. S4** Timelapse montage of fluorescence response to 40 mA electrical stimulation in labeled mouse hippocampus (scale: 100 μm).

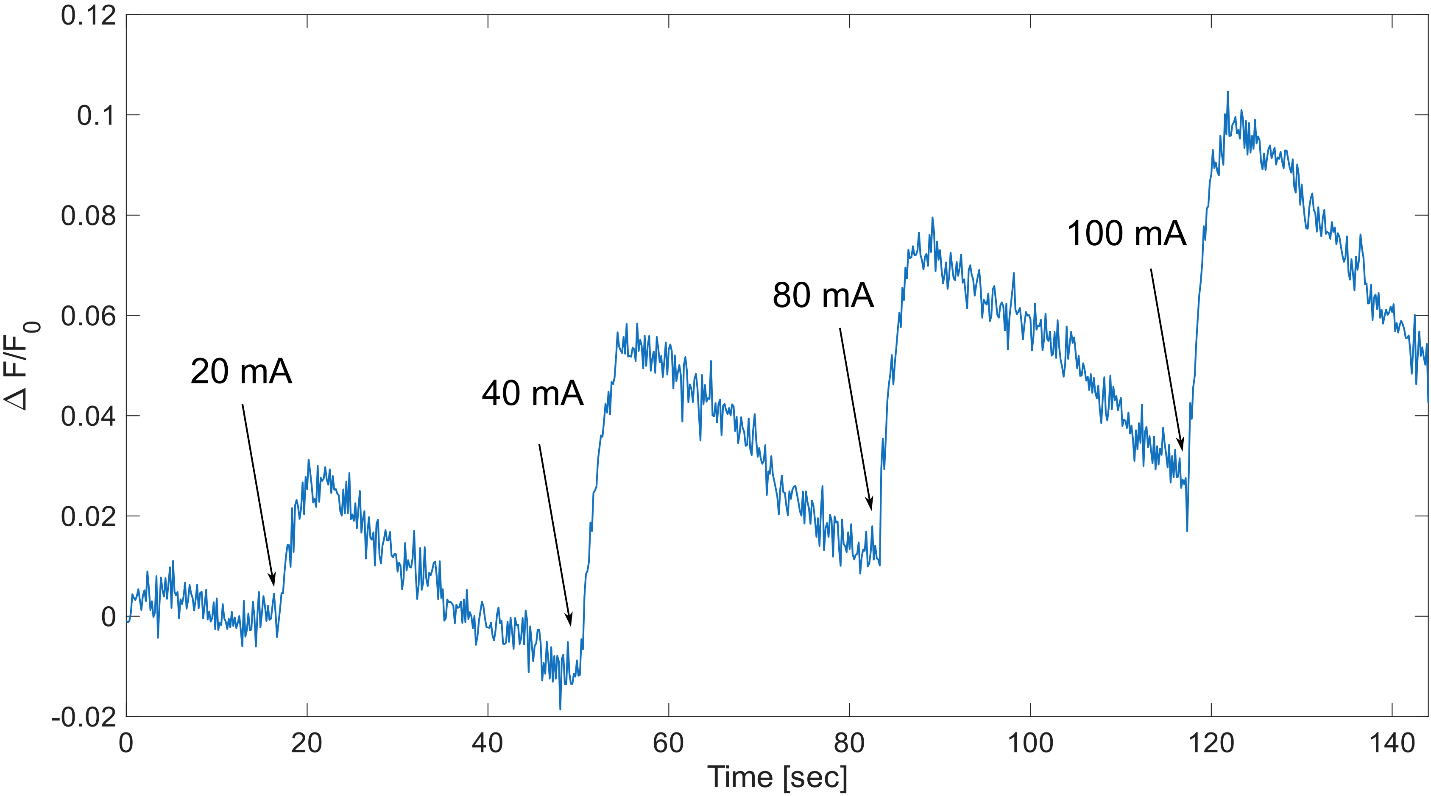

**Fig. S5** Fluorescence intensity responses to repeated electrical stimulations of progressively higher amperages.

**Fig. 1** Optical components and molecular design of the DNA-aptamer potassium sensor.

**Fig. 2** Optical properties of the DNA-aptamer potassium sensor in solution.

**Fig. 3** K^+^ sensor labeling of cell and tissue models.

**Fig. 4** Electrical stimulation of labeled acute brain slices.
